# Conformational changes linked to ADP release from human cardiac myosin bound to actin-tropomyosin

**DOI:** 10.1101/2022.09.23.509077

**Authors:** M.H. Doran, M.J. Rynkiewicz, D. Rassici, S.M.L. Bodt, M. E. Barry, E. Bullitt, C.M. Yengo, J.R. Moore, W. Lehman

## Abstract

Following binding to the thin filament, β-cardiac myosin couples ATP-hydrolysis to conformational rearrangements in the myosin motor that drive myofilament sliding and cardiac ventricular contraction. However, key features of the cardiac-specific actin-myosin interaction remain uncertain, including the structural effect of ADP release from myosin, which is rate-limiting during force generation. In fact, ADP release slows under experimental load or in the intact heart due to the afterload, thereby adjusting cardiac muscle power output to meet physiological demands. To further elucidate the structural basis of this fundamental process, we used a combination of cryo-EM reconstruction methodologies to determine structures of the human cardiac actin-myosin-tropomyosin filament complex at better than 3.4 Å-resolution in the presence and in the absence of Mg^2+^·ADP. Focused refinements of the myosin motor head and its essential light chains in these reconstructions reveal that small changes in the active site are coupled to significant rigid body movements of the myosin converter domain and a 16-degree lever arm swing. Our structures provide a mechanistic framework to understand the effect of ADP binding and release on human cardiac β-myosin and offer insights into the force-sensing mechanism displayed by the cardiac myosin motor.

**Short Summary:** Cryo-EM was used to elucidate high-resolution structures of actin-tropomyosin filaments decorated with human cardiac myosin in the rigor and ADP-bound states. Differences in the myosin lever arm orientation detected correlate with the overall actomyosin-linked force-sensitivity.

## Introduction

Myosin motors are part of a large protein superfamily that bind to and generate force on actin tracks(Coluccio, 2020). In striated muscle cells thick filaments, myosin motor heads project laterally from the filament axes, while their tails form the filament shaft. Here, in a cyclic fashion, the motor heads bind to and release from thin filament actin, powering the sliding of thin filaments past thick filaments(Huxley, 1969; Lymn and Taylor, 1971). As part of this process, actin-binding catalyzes the release of ATP hydrolysis products, thus coupling the actomyosin ATPase cycle to force generation, i.e. transducing chemical energy into mechanical work(Geeves and Holmes, 2005; Geeves, 2016).

The structure of all myosin motor heads is made up of four subdomains, including the upper and lower 50 kDa domains (U50, L50), an N-terminal domain, and a converter domain (see Figure 1A). The actin-binding region sits at the tip of the motor domain and extends over a wide region between the upper and lower 50K domains. At the rear of the motor head, a long α-helix “lever-arm” connects to the converter domain and, in muscle thick filaments, bridges between the motor head the filament shaft. Connecting these subdomains are a series of myosin surface loops referred to as loops 1, 2, 3, 4, the cardiomyopathy loop, and the activation loop.

**Figure 1:**
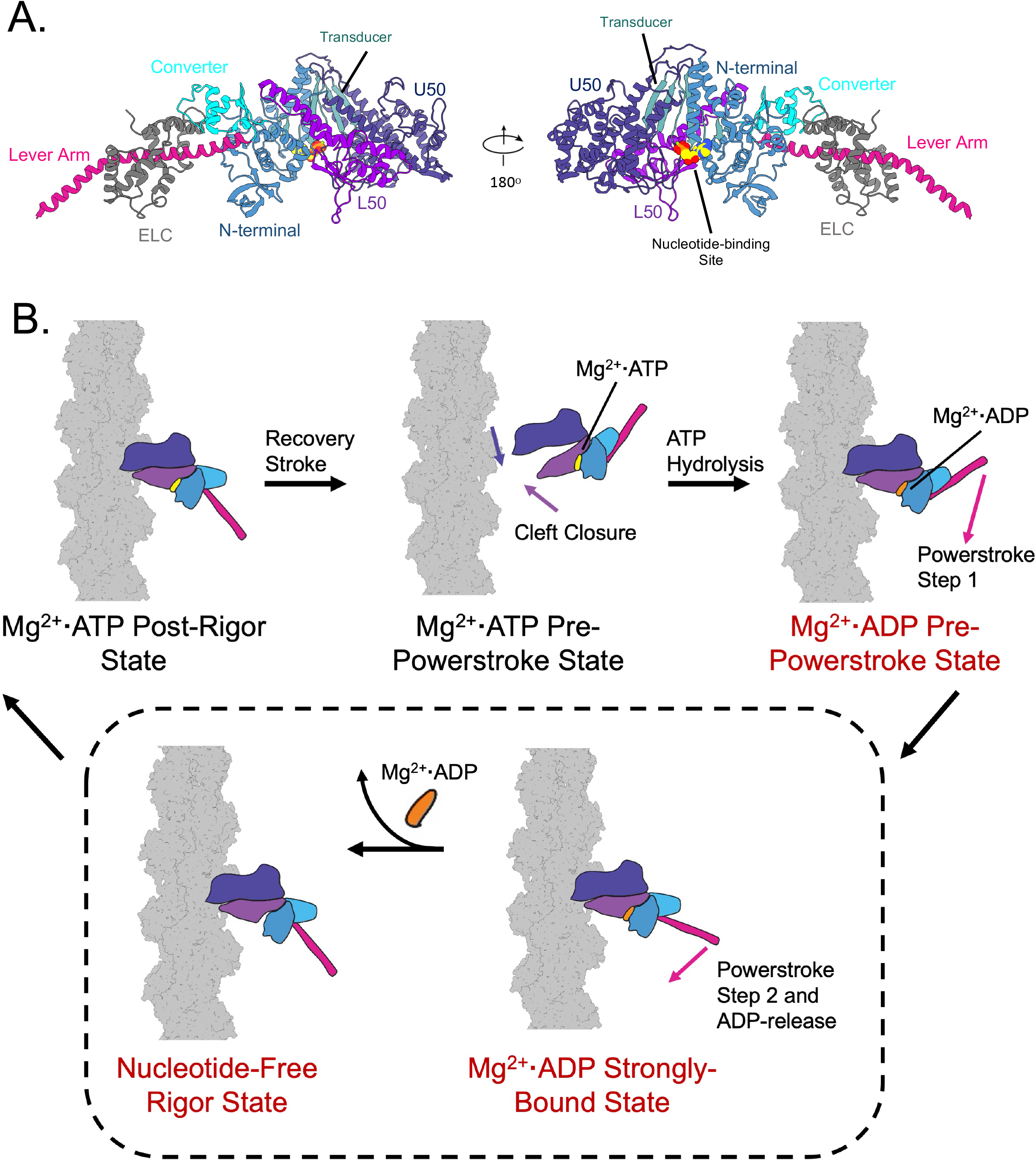
Conformational changes in the myosin motor domain produce force on actin. Structural organization of the myosin motor. A). Ribbon diagram of the myosin S1 motor domain displayed in two orientations. The secondary structure of the molecule is colored according to the subdomains of the protein, including the upper 50 kDa domain (U50, dark purple), the lower 50 kDa domain (L50, light purple), the N-terminal domain (blue), converter (cyan), lever arm (magenta), and the transducer region (light blue). The nucleotide binding site is marked by a space filling model of ADP/Mg. This figure was adapted from (31) and uses the coordinated deposited in the PDB: 5H53. B). Schematic of the myosin force generating mechanism. β-cardiac myosin II undergoes conformational changes that are paired to actin binding and nucleotide-hydrolysis product release steps in order to generate the requisite force that results in cardiac muscle contraction. This figure describes each step of the cycle and the rearrangements of the myosin head, including cleft closure and the movement of the lever arm during the powerstroke. Each subdomain of the molecule is colored as in Figure 1A, while the force generating steps are shaded red. In order to provide clarity, the phosphate release step, which occurs after hydrolysis and before the first step of the powerstroke, was not included in this figure. The step we focus on in this paper, ADP-release, is outlined in the dashed box.

The nucleotide binding site lies between the U50 and the N-terminal domain above a seven-stranded beta sheet “transducer” domain (see Figure 1A). The transducer, in turn, links actin-associating regions to the nucleotide binding site, and is thought to transduce nucleotide binding to structural changes that control myosin attachment and detachment from actin(Coureux et al., 2004; Gurel et al., 2017). Moreover, shifts in the converter domain cause the lever arm to swing, generating significant force, a process commonly referred to as the powerstroke. Thus, the repeated attachment and detachment of myosin heads on actin is the molecular basis of the crossbridge cycle on F-actin and the corresponding sliding of filaments during muscle shortening (Geeves and Conibear, 1995).

Although some details remain controversial, the basic outlines of the crossbridge cycle are generally accepted (shown in Figure 1B). This cycle consists of ATP hydrolysis followed by product release steps that are paired with conformational changes within the motor. When bound to Mg^2+^**·**ATP, myosin displays demonstrably low actin-affinity in a “pre-powerstroke-state” conformation (PPS). In the PPS conformation, the motor domain lever arm is primed in an up position and the cleft between U50 and L50 is open. Here, myosin hydrolyzes ATP rapidly and the resulting myosin-ADP-P_i_ complex initially binds to actin via weak initial electrostatic interactions(Furch et al., 2000; Joel et al., 2001; Onishi et al., 2006). To continue the cycle, several events must occur, including Pi release, U50/L50 cleft closure, and lever arm rotation of the powerstroke. Although these events are tightly coupled, the timing of each remains debated (Llinas et al., 2015; Tang et al., 2021; Moretto et al., 2022; Debold, 2021). Nevertheless, during U50/L50 cleft closure, the upper and lower 50K domains rotate towards one another, resulting in stronger binding to F-actin(Milligan et al., 1990).

In addition to cleft closure and strong actin binding, the myosin lever arm rotates and undergoes the powerstroke movement, leaving the Mg^2+^**·**ADP-associated myosin (Houdusse and Sweeney, 2016). Finally, Mg^2+^**·**ADP is released from myosin, which is thought to initiate a second, smaller rotation of the lever arm in the β-cardiac motor. This last step, which is rate-limiting for the shortening velocity of cardiac muscle, has been shown to be strain-dependent and myosin isoform-specific(Siemankowski and White, 1984; Sweeney and Houdusse, 2010; Weiss et al., 2001). The isoform-specific conformational changes associated with the step may be opposed by load, resulting in a slower release of Mg^2+^**·**ADP from myosin and myosin from actin(Cremo and Geeves, 1998). After nucleotide-release, myosin is left in the nucleotide-free “rigor-state” state, where subsequent ATP binding initiates actin dissociation and a new cycle.

Cryo-EM methods have been used to generate reconstructions of many actomyosin complexes, including ones for mammalian bovine and porcine cardiac myosin(Doran et al., 2020; Risi et al., 2021). However, only three high-resolution cryo-EM solutions have been solved for ADP-associated actomyosin, all containing non-sarcomeric myosins (containing myosin V, myosin IB, and myosin 15) and hence none including striated muscle myosin isoforms(Pospich et al., 2021; Gong et al., 2022; Mentes et al., 2018). In the current study, we focused on correcting this deficit by reconstructing high-resolution structures of F-actin decorated with mammalian cell expressed β-cardiac myosin II under nucleotide-free conditions or in the presence of Mg^2+^**·**ADP. Notably, β-myosin is the major motor protein found in cardiac ventricular muscle and is also well-recognized as the site of over 300 pathogenic mutations that lead to hypertrophic (HCM) and dilated (DCM) cardiomyopathy(Morimoto, 2007; Homburger et al., 2016).

The cryo-EM reconstructions presented in this paper contain actin-tropomyosin filaments associated with nucleotide-free and ADP-bound human β-cardiac myosin II. The coordination of Mg^2+^**·**ADP within the myosin active site as well as the conformation of the nucleotide-free pocket are well-defined. In addition, comparison of the two structures elucidates the conformational changes associated with ADP release from myosin. We observe the previously described opening of the active site, which includes the movement of the HF helix and surrounding loops away from the nucleotide. In addition, we reveal the isoform-specific differences in the other parts of the motor domain, namely the 3 Å difference of the relay helix, the 6.5 Å alteration of the converter domain and the 16-degree rotation of the lever arm. Overall, our structures provide a roadmap to determine how disease-causing mutations disrupt the cardiac myosin ADP-bound-to-rigor transition.

### Materials and Methods Protein Purification

Porcine cardiac F-actin, identical in sequence to human cardiac α-actin, and human cardiac α-tropomyosin, containing an N-terminal Ala-Ser extension, were prepared as previously described(Sundar et al., 2020). Recombinant human β-cardiac myosin S1, including residues 1-842 and containing C-terminal Avi and FLAG tags, was expressed in C2C12 mammalian cells and purified with FLAG affinity chromatography(Tang et al., 2019; Swenson et al., 2017). The initial recombinant adenovirus was purchased from Vector Biolabs (Malvern, PA) and amplified to generate high-titer stocks for infection of C2C12 cells (Touma et al., 2022; Liu et al., 2015). The purified myosin contained mouse skeletal muscle regulatory and essential light chains(Swenson et al., 2017).

### Sample preparation for cryo-EM

Freshly glow-discharged R1.2/1.3 Quantifoil gold grids, which were discharged with a Pelco EasiGlow (Electron Microscopy Sciences, Hatfield, PA) were used in our work. Actin-tropomyosin filaments were prepared by mixing 3 μM porcine cardiac actin and 9 μM human cardiac tropomyosin in 10 mM sodium acetate, 3 mM MgCl_2_, 1 mM DTT and 10 mM MOPS buffer, at pH 7.0. Directly before adding this sample to the grid, 0.3% of the surfactant octyl β-D-glucopyranoside (Sigma-Aldrich, St. Louis, MO) and 0.1% bacitracin were added to the actin-tropomyosin mixture. These additives facilitated even and thin ice as well as maximized filament spreading. 1.5 μl of the actin-tropomyosin solution was applied to the gold grid and manually blotted for 1 s at 10 ° C and 100% humidity. Next, for the nucleotide-free complex, 1.5 μL of solution containing 9.5 μM myosin S1 and 0.3% of octyl β-D-glucopyranoside was applied to the grid, blotted for 6 s, and immediately plunge frozen in liquid ethane using a Vitrobot Mark III system (FEI/Thermo Fisher Scientific, Hillsboro, OR) as previously described(Doran et al., 2020). For the complex containing Mg^2+^**·**ADP, the same process was repeated, except 10 mM MgCl_2_ and 1 mM ADP were added to the myosin solution.

### Cryo-EM data collection

Samples were examined with a Titan Krios transmission electron microscope operating at 300 kV (Thermo Fisher Scientific, Hillsboro, OR) at the Purdue Cryo-EM Facility (West Lafayette, IN). Cryo-EM movies were collected on a Gatan K3 Summit direct electron detector (Gatan/AMETEK, Berwyn, PA) using Leginon data acquisition software (Suloway et al., 2005) at a nominal magnification of 81,000 x corresponding to a pixel size of 0.539 Å using a 20 eV energy filter and with a dose rate of 17.21 e^-^/Å^2^/s. We used a total exposure of 3.12 seconds, which resulted in an accumulated dose of 53.70 e^-^/Å^2^. Frames were recorded every 0.08 seconds for a total of 40 frames per micrograph. 3,909 movies were collected for the actin-tropomyosin-myosin complex and 3,961 were collected for the actin-tropomyosin-myosin-ADP complex.

### Helical Reconstruction

In order to solve the structure of the human cardiac myosin bound to actin-tropomyosin in the presence and absence of ADP, we used helical reconstruction methods as implemented in RELION 3.1.1 (Supporting Material Figures 1 and 2). Micrographs were first motion corrected using MotionCor2 (Zheng et al., 2017) using 5×5 patches and binning pixels 2x for a final size of 1.078 Å. After motion correction, the contrast transfer function (CTF) was estimated using CtfFind 4.1.13(Rohou and Grigorieff, 2015). Initially, filaments from ∼200 micrographs were manually selected and divided into overlapping segments (47 nm long, 440 pixels) with an intersegment displacement of 27.5 Å. Reference-free 2D classification was then performed to generate templates for autopicking. After autopicking, 2D classification was performed in order to remove false positives as well as ice-contaminated and damaged segments. Segments within 2D classes that contained secondary structure features were selected for 3D classification with 5 classes using a featureless cylinder as a reference. Initial helical parameters of -166.5 degrees and 27.5 Å were used. The 131,194 segments (ADP-bound) and 176,178 segments (rigor) contributing to the classes with the highest resolution were selected and used for subsequent refinements. In the first round of 3D auto-refinement, a featureless cylinder was again used as a reference to produce our initial reconstructions. During this auto-refine job, we also performed local helical parameter searches. For both samples, our twist search ranged from -166.2 degrees to -167.6 degrees, while our rise search ranged from 27.2 Å to 27.9 Å. Both the rigor and nucleotide-free reconstructions converged on a twist of -166.8 and a rise of 27.9 Å. After postprocessing in RELION, these reconstructions reached resolutions of 4.0 Å for the rigor complex and 4.3 Å for the ADP-bound complex as calculated using the FSC_.143_ criterion.

After the initial reconstructions, we sought to increase the resolutions of the helical reconstructions. To do this, we performed two iterations of CTF refinement, Bayesian polishing, and auto-refine. For each CTF refinement, the B-factor estimation option was not used as it led to spurious resolution estimates and distorted maps. For each auto-refine, the reconstruction from the last refinement, low-pass filtered to 30 Å was used as the initial model. Additionally, for each of these auto-refine jobs, we imposed a solvent mask that included the central 6 actin-subunits, a single tropomyosin dimer, and 3 myosin heads, which improved the resolution and quality in these regions of the reconstruction dramatically. A third iteration of CTF refinement, polishing, and auto-refine did not improve the resolution. Final postprocessing of the second iteration 3D auto-refine using a 45% Z length mask yielded our final maps. Using the FSC_.143_ criteria, the final resolutions were reported to be 3.4 Å for nucleotide-free and 3.3 Å for myosin ADP-bound filaments. Local resolution measurements of the maps showed a resolution range of between 3.2 (near the central axis of the filament) and 10 Å (for the outer edge of myosin), thus poor density was noted for the lever arm/essential light chain region of the myosin.

### Single-Particle Reconstruction of the Myosin/Light Chain Complex

Single-particle, focused refinement was performed on individual myosin heads from our helical reconstructions in order to improve resolution of regions of S1 at high radius from the actin filament in our reconstructions, (see Supporting Material Figures 1 and 2). To do this, we first created masks that enveloped the central S1 head of the helical reconstruction to focus the classifications/refinements solely on that region. To create these masks, we rigid body docked the nucleotide-free skeletal muscle myosin-essential light chain structure (PDB ID:5H53) into the helical reconstruction of the nucleotide-free structure using the fit-in-map tool in UCSF chimera. To generate a mask for the ADP-bound form, the 5H53 structure was rigid body fit into the ADP form of the reconstruction density. Because 5H53 contained a nucleotide-free conformation, the lever arm and light chain did not fit within the density. To fit them to the helical density, we performed a single round of refinement using Phenix. After fitting to the helical reconstructions, the PDB files were converted into a volume using the eman2 command, e2pdb2mrc.py and then into a soft mask in RELION with a 5-pixel extension and a 5-pixel soft-edge.

A single round of 3D classification was carried out using these masks without alignment or imposed helical restraints on our final helical reconstruction particle sets in order to select data that contained recognizable lever arm and light chain density. After picking the classes that contained secondary structural elements in these regions, we were left with final data sets with 88,048 particles of nucleotide free myosin and the ADP-bound class contained 107,605 particles of myosin with bound ADP. Following classification, which identified the segments containing the high-quality density for the outer regions of the myosin, we performed a single round of 3D auto-refine without imposing helical symmetry on the selected segments. These reconstructions were completed using the final helical reconstructions, low-pass filtered to 4 Å, as references while performing only 1.8 degrees of local angular searches with our central myosin solvent masks. The 3D auto-refine jobs produced myosin motor domain reconstructions with greatly improved density at the active site, lever arm, converter domain, and the essential light chain (see Supporting Material Figure 4). Final postprocessing in RELION resulted in resolutions of 3.83 for the rigor complex and 3.79 Å for the ADP-bound complex (Supporting Figure 3). The primary maps used for model building in the single-particle reconstructions were low pass filtered to 3.85 Å during post-processing in order to reduce the amount of noise in the reconstruction and prevent overfitting of the model.

### Model Building

To start, we first modelled the rigor and ADP-bound complexes into our helical reconstructions. The models based on the helical reconstructions will be referred to as the rigor-HA and ADP-HA, respectively. After completing these models, we then used our single-particle reconstructions to build into the improved density of the myosin S1 and essential light chain portions of the complex. The models built into the single-particle reconstructions will be called rigor-SP and ADP-SP respectively.

Our starting structures for the rigor-HA and ADP-HA models consisted of the central actin chain from the Yamada et al. high-calcium thin filament reconstruction (chain G from PDB ID:6KN8) and myosin and tropomyosin chains from the bovine rigor structure (chains G and chains O,P respectively from PDB ID:6X5Z), which were rigid body fit in Chimera into the helical reconstructions. The human myosin structure was threaded onto the bovine coordinates before fitting using SWISS-MODELLER(Schwede et al., 2003). After initial fitting, the model was subjected to iterative rounds of manual rebuilding in Coot and refinement in Phenix. Each refinement cycle consisted of conjugate gradient minimization, rigid body fitting (using all atoms as a single rigid body), and local grid search in Phenix. To maintain good geometry, secondary structure and Ramachandran restraints as well as ideal bond and angle restraints (target rmsd of 0.005 Å and 0.5 degrees, respectively) were applied. Portions of the model that fell outside of clear density were not included. The final model of the rigor form, for example, consisted of actin residues 5-375, myosin residues 6-199, 216-624, 634-728, and 738-780, and tropomyosin residues 47-210 (pseudorepeats 3-5 as used for 6X5Z). Progress of the refinement was monitored using the cryo-EM validation tools in Phenix, in particular, improvements to the clash score were checked at each round. For the ADP-HA model, to include the nucleotide, we manually added the Mg^2+^·ADP atoms into the density surrounding the HF helix using the pdb: 6C1H as an initial guide. After initial rigid-body fitting in Chimera, we then manually refined its position in Coot 0.9.6 to prepare for the Phenix refinements.

After building into the helical reconstructions, we proceeded to build into the single-particle maps to extend our models at the lever arm and essential light chain. For both the rigor-SP and ADP-SP forms, we fit the myosin motor domains into the single-particle reconstruction density in Chimera. Once fitted to the map, we manually added residue-by-residue to the lever arm using Coot, extending out to residue 796, where clear density ceased. After extending the lever arm, we then modelled the light chain using an AlphaFold generated pdb of the murine Myl1 gene(Jumper et al., 2021). First, the AlphaFold pdb, spanning residues 43-188, was rigid-body docked into the single-particle density using Chimera. Although the secondary structure fit well, the N-terminus of the molecule (1-42) was not included because our reconstruction lacked density for this region. After fitting the complex, the myosin-essential light chain model was subjected to iterative rounds of manual rebuilding in Coot and refinement in Phenix as was done when building into the helical models. Validation statistics for each of the models can be found in Table 1: Cryo-EM data collection, refinement, and validation statistics.

**Table 1:** Cryo-EM data collection, refinement, and validation statistics. Compilation of the relevant statistics for the cryo-EM reconstructions and subsequent atomic models.

|  | Rigor-HA | ADP-HA | Rigor-SP | ADP-SP |
| --- | --- | --- | --- | --- |
| **Data collection and processing** |  |  |  |  |
| Magnification | 81,000x | 81,000x | 81,000x | 81,000x |
| Voltage (kV) | 300 | 300 | 300 | 300 |
| Electron exposure (e-/Å<sup>2</sup>) | 53.62 | 53.62 | 53.62 | 53.62 |
| Defocus range (μm) | -1.0 ~ -3.0 | -1.0 ~ -3.0 | -1.0 ~ -3.0 | -1.0 ~ -3.0 |
| Pixel size (Å) | 1.078 | 1.078 | 1.078 | 1.078 |
| Symmetry imposed | Helical Symmetry | Helical Symmetry | None | None |
| Initial segment images (no.) | 1,045,903 | 628,787 | 176,178 | 131,194 |
| Final segment images (no.) | 176,178 | 131,194 | 88,048 | 68,358 |
| Map resolution (Å) | 3.4 | 3.3 | 3.8 | 3.8 |
| FSC threshold | 0.143 | 0.143 | 0.143 | 0.143 |
| Map resolution range (Å) | 3.3-7.3 | 3.2-10.5 | 3.8-4.8 | 3.5-6.1 |
| **Refinement** |  |  |  |  |
| Initial model used (PDB code) | 6X5Z | 6X5Z | Rigor-HA | Rigor-HA |
| Model resolution (Å) | 3.5 | 3.5 | 4.4 | 4.1 |
| FSC threshold | 0.5 | 0.5 | 0.5 | 0.5 |
| Model resolution range (Å) | 3.1-3.5 | 3.0-3.4 | 3.6-4.4 | 3.4-4.1 |
| Map sharpening *B* factor (Å<sup>2</sup>) | -71.49 | -52.13 | -83.37 | -72.94 |
| Model composition |  |  |  |  |
| Non-hydrogen atoms | 23,281 | 23,254 | 7,107 | 7,135 |
| Protein residues | 2929 | 2921 | 886 | 886 |
| Ligands | 5 | 6 | 0 | 1 |
| *B* factors (Å<sup>2</sup>) |  |  |  |  |
| Protein | 112.33 | 85.14 | 203.27 | 218.65 |
| Ligand | 69.31 | 54.89 | N/A | 151.65 |
| R.m.s. deviations |  |  |  |  |
| Bond lengths (Å) | 0.003 | 0.002 | 0.002 | 0.002 |
| Bond angles (°) | 0.575 | 0.544 | 0.492 | 0.515 |
| Validation |  |  |  |  |
| MolProbity score | 1.84 | 2.29 | 1.70 | 1.70 |
| Clashscore | 7.28 | 6.93 | 8.25 | 8.23 |
| Poor rotamers (%) | 0.20 | 0.20 | 0.13 | 0.52 |
| Ramachandran plot |  |  |  |  |
| Favored (%) | 97.04 | 96.45 | 96.23 | 96.23 |
| Allowed (%) | 2.96 | 3.55 | 3.77 | 3.77 |
| Disallowed (%) | 0.00 | 0.00 | 0.0 | 0.0 |

## Results and Discussion

A major goal of our ongoing studies is to determine cryo-EM structures of samples containing human myofilament constituents in order to define the structure of the human myosin motor head and its interface with F-actin-tropomyosin. In the current work, we focused on solving the structure of the myosin head when it is strongly bound to thin filaments in the presence and absence of Mg^2+^**·**ADP. Our samples contained human β-cardiac myosin subfragment 1 (S1) engineered in C2C12 mammalian cells, which encompasses the myosin actin-binding region, its nucleotide binding, and catalytic site as well as its mechano-sensitive converter, force-producing lever arm, and light chain elements. We previously showed that the expressed S1 displays native motor and enzymatic activity(Swenson et al., 2017). Since the myosin S1 lacks the myosin tail it does not form filaments but saturates binding sites along actin-tropomyosin filaments in our reconstructions.

We prepared samples of S1-decorated actin-tropomyosin filaments in the absence and in the presence of Mg^2+^**·**ADP, recorded cryo-EM images, and solved respective cryo-EM structures using a two-step approach. Here, we first applied helical averaging methods in RELION to obtain 3.3 -3.4 Å resolution for the central portion of the reconstructions and then employed single particle, focused refinement methods specifically directed at the S1 densities to obtain improved resolution and quality for parts of myosin at a higher radius from the actin filament axis. In this way, we achieved high-quality maps over the entire structure including the distal myosin lever arm and associated essential light chain. The results of the two procedures were combined to produce composite maps with high quality density over the entire complex, as shown in Figure 2. Once elucidated, these cryo-EM structures were used to evaluate the transition between force producing states of myosin strongly bound to actin.

**Figure 2:**
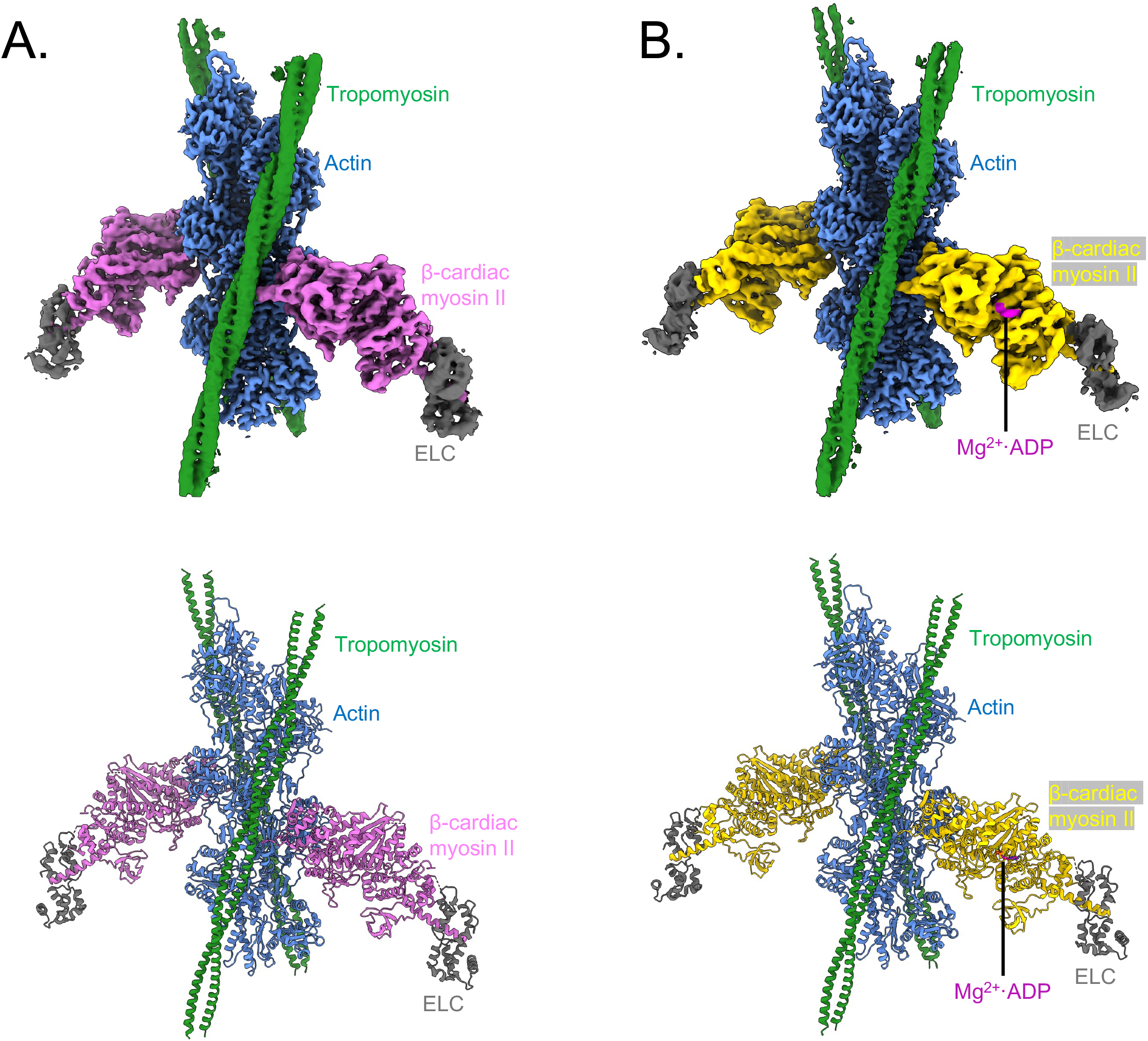
Composite cryo-EM reconstructions of the human cardiac actin-β-myosin-tropomyosin complex. A). Top panel: composite cryo-EM map of the rigor actin-β-cardiac myosin II-tropomyosin complex. These reconstructions were created by combining the central portion of the actin-tropomyosin from the helical reconstructions and the S1 domain of myosin, including the essential light chain (ELC) from focused refinements. This figure was rendered in Chimerax using the surface zone tool (Pettersen et al., 2021). Bottom panel: atomic model of the complex in the same orientation as the reconstruction above. B). Top panel: Mg^2+^**·**ADP-bound complex composite map. The rendering of the ADP-bound complex was completed in the same fashion as the rigor complex. Bottom panel: atomic model of the rigor complex in the same orientation as the reconstruction. The nucleotide is also indicated in both the reconstruction and model.

### High resolution cryo-EM reconstructions of the rigor and ADP-bound complexes

Micrographs of F-actin-tropomyosin decorated with S1 displayed the typical arrowhead pattern in both nucleotide-free and in Mg^2+^·ADP-containing buffer (Supporting Figures 1 and 2). As mentioned, our initial helical reconstructions of the fully-decorated filaments yielded global resolutions of 3.3 Å and 3.4 Å for the actomyosin-tropomyosin filaments in the presence or absence of Mg^2+^**·**ADP respectively. Despite the high reported resolution, the local resolution varied. The resolution was highest in the interior of the actin and at the actin-myosin interface, but decreased radially from the actin filament axis (Supporting Figure 3). Because of this distribution, the resolution of the helical reconstructions is the lowest at the outer edge of the myosin motor domain and only poor density at low contour is observed for the essential light chain region. Following application of focused single-particle refinement of this region, the motor domain, with the lever arm, and associated essential light chain, reached a global resolution of 3.8 Å. Thus, the resolution of the full cardiac actomyosin complex (Figure 2) is improved over that achieved previously for striated-muscle isoforms(Doran et al., 2020; Risi et al., 2021).

Our previously solved nucleotide-free bovine masseter β-myosin structure fits extremely well into the human cardiac model reconstruction and was used as an initial reference to build new refined structures based on the human cardiac myosin sequence(Doran et al., 2020). In our helical reconstructions, most side-chains are evident for the actin filament and the myosin motor domain at the actin-myosin interface. Following single-particle analysis, the motor domain itself is also well-defined and large amino acid side chains are visible. When present, Mg^2+^**·**ADP density is obvious, allowing identification of corresponding coordinating side chains as shown in Figure 3. In distal regions of the motor, including the converter domain and the lever arm, secondary structure is visible in helical reconstructions and resolved definitively out to myosin residue 796 in our composite maps that incorporate the single particle refinements. The tropomyosin coiled-coil is also clearly visible in these reconstructions traversing actin subdomains 3 and 4 of successive actin protomers as previously described(von der Ecken et al., 2016; Doran et al., 2020; Risi et al., 2021), but residue-level detail is not resolved due to helical averaging(Milligan et al., 1990; Vibert et al., 1997).

**Figure 3:**
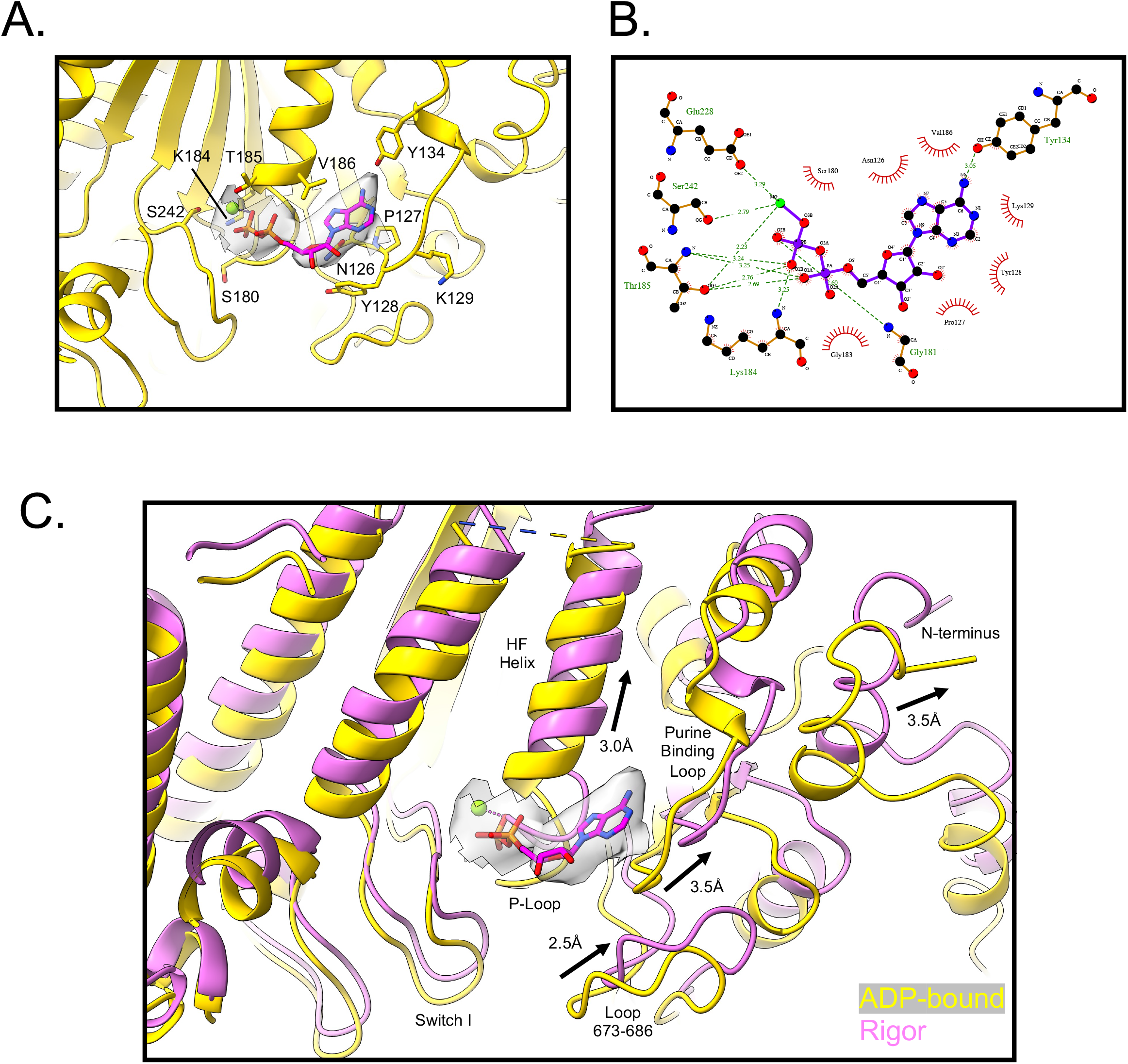
Coordination of the ADP-Mg^2+^ and active site differences between ADP-bound and rigor. A). View of the nucleotide-binding pocket and the placement of the Mg^2+^**·**ADP within it. The density of the nucleotide from the single particle focused refinement is shown in grey. Side chains that are involved in the coordination and binding of the nucleotide are displayed around the ADP. B). Ligplot diagram detailing the interaction between the myosin active site and nucleotide. Hydrogen bonds are shown in green, while hydrophobic contacts are shown by red semi-circles. The orientation of the nucleotide is similar to 3A. C). Comparison of the active site in the rigor complex (orchid) vs. the Mg^2+^**·**ADP-bound complex (gold). The changes of each part of the nucleotide pocket are displayed including the movement of the HF helix and P-loop, switch I, the purine-binding loop, the N-terminus, and loop 673-686. These comparisons were created by aligning the actin filament of the helical models and overlaying the single particle S1 models on their helical counterparts using the matchmaker command in ChimeraX.

Our two reconstructions are our surrogates for the β-cardiac myosin’s ADP-bound and free states when tightly bound to actin. Comparison of these structures, as shown in Figures 3 and 4, is the focus of our study and reveals ADP-dependent reconfigurations. However, at the actomyosin interface, the models demonstrate remarkable similarity and the regions are nearly superimposable. In both structures, the cleft separating upper and lower 50K domains is closed and the helix-loop-helix motif and cardiomyopathy loop tether the motor to the actin filament mainly via hydrophobic contacts. The myosin surface loops 2, 3, 4, and the activation loop form a secondary interface, without significant differences from previous work (Doran and Lehman, 2021). The conservation of the actomyosin interface in the ADP-bound and rigor states coincides with actin-binding affinity measurements, which report that both states bind to F-actin with nanomolar affinities(Yengo et al., 2002).

**Figure 4:**
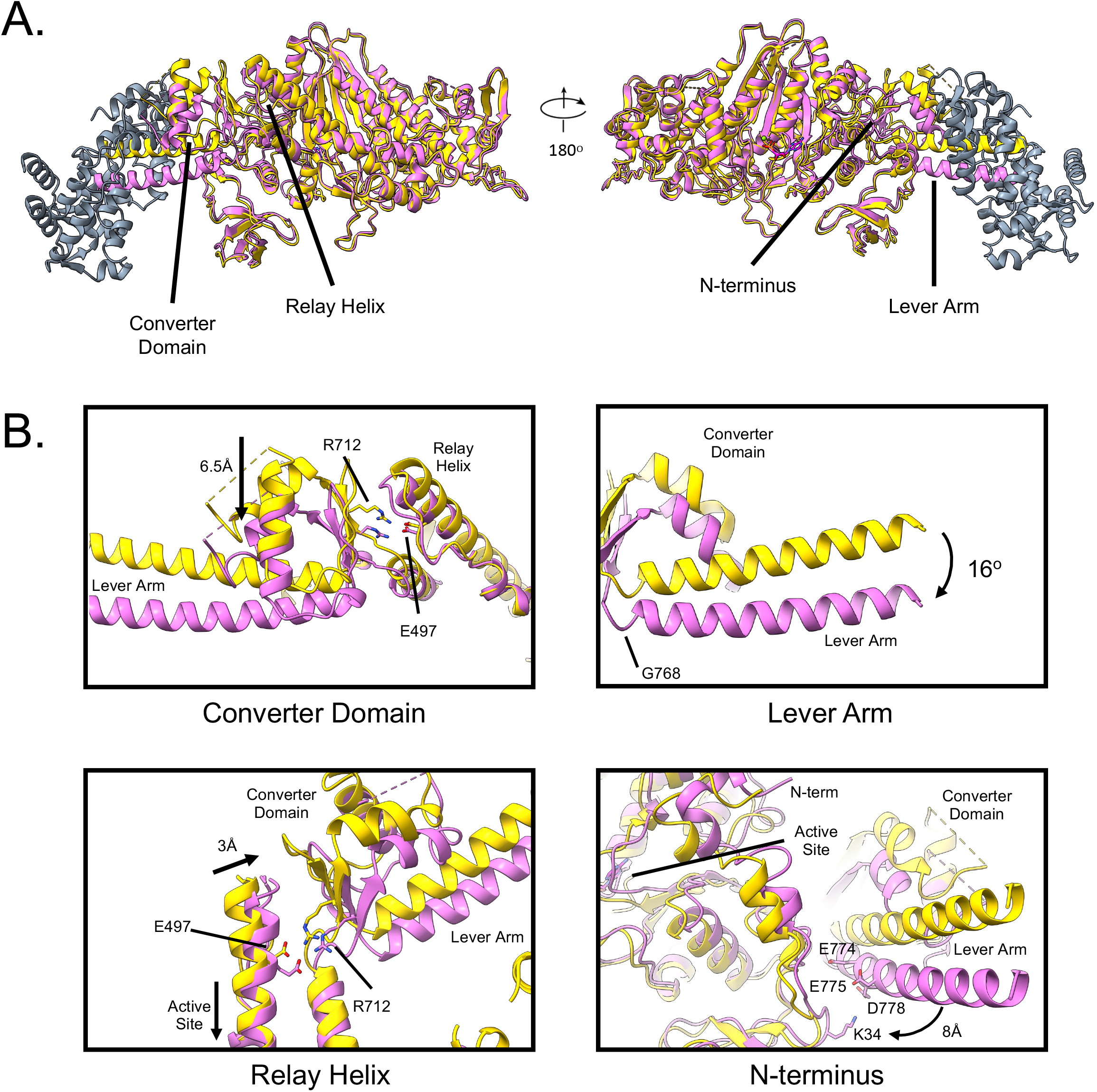
Conformational changes in the motor domain associated with ADP-Mg^2+^ release. Comparisons of the nucleotide-free rigor form and Mg^2+^**·**ADP-bound complex reveal conformational changes associated with ADP release. A). Comparison of the S1 motor domains of the nucleotide free-rigor form (orchid) and the Mg^2+^**·**ADP-bound complex are shown in two different orientations. We highlight significant portions of the motor domain that change from between the models including the converter domain, relay helix, N-terminus, and the lever arm. B) Individual panels show the conformational differences in more detail for the specified regions. Small differences at the active site are linked to these larger movements of subdomains. The converter domain change was measured from the α-carbon of residue S747. The lever arm rotation was measured using the axes, planes, and ellipsoids tool in Chimera. The axes used in the calculation was defined as the last 13-resiudes of the lever arm. The relay helix change was measured by the change in the α-carbon of E504.

### Cryo-EM reveals the coordination network of Mg^2+^·ADP in the myosin active site

In the Mg^2+^**·**ADP-bound reconstruction, the nucleotide can be easily fitted to densities in the myosin active site and the coordinating residues associated with the nucleotide are clearly identified (see Figure 3). The coordination of Mg^2+^**·**ADP at the active site (Figure 3) contains a Walker A motif (P-loop) that coordinates the β-phosphate of the ADP using the positively charged residue K184, while T185 and E228 help coordinate the magnesium ion(Wendler et al., 2012; delToro et al., 2016; Walker et al., 1982).

Apart from the P-loop, polar and non-polar residues from the surrounding purine-binding loop (residues 126-135), HF helix (residues 184-197), as well as switch I, surround the nucleotide to form a tight coordination network with the Mg^2+^**·**ADP (Figure 3A, B). Here, Y134 of the purine binding loop forms a hydrogen bond with the adenine ring, while the residues, Y128 and P127 form hydrophobic contacts. The backbone nitrogen atoms of HF helix residues G183 and G181 form hydrogen bonds with the ADP phosphate groups. Additional contacts with ADP phosphates are formed by switch I, particularly residue S242, which interacts with the accompanying magnesium adjacent to the β-phosphate of ADP. The ADP-coordination described above closely matches the active site in the crystal structure of the actin-free “post-rigor” bovine β-cardiac myosin in complex Mg^2+^**·**ADP (Robert-Paganin et al., 2018). Comparing these active site architectures with published coordinates of other Mg^2+^**·**ADP complexed actomyosin isoforms, such as myosin V, 1b, and 15, we find that they are nearly superimposable, highlighting the evolutionary conservation of this part of the myosin structure (Mentes et al., 2018; Pospich et al., 2021; Gong et al., 2022).

### Comparisons of the ADP-bound and nucleotide free active sites show changes at the active site

Comparing our reconstructions in the Mg^2+^**·**ADP complex with the rigor complex show the structural changes in the active site associated with release of Mg^2+^ and ADP from myosin (Figure 3C). First, this entails a 3.0Å change in HF helix position away from the nucleotide binding site (as measured by the α-carbon for residue V186). This change would cause T185 to lose coordination with Mg^2+^, which could destabilize nucleotide-binding. In fact, previous studies have identified the magnesium ion as an essential regulator for myosin ADP-release(Rosenfeld et al., 2005; Swenson et al., 2014). Beyond the movement of the HF helix, when comparing the ADP-bound and rigor form, differences are also found in the 7 stranded β-sheet transducer, which, as mentioned in the introduction, is thought to couple active site changes to rearrangements in other parts of the motor(Gurel et al., 2017). Here, it appears that the transducer modestly twists between the two states, leading to an alteration in the interaction network between the two forms. The twist of the transducer therefore leads to alterations in the active site coordination that could aid in nucleotide-release.

Additional changes also occur in the loops surrounding the HF-helix. For instance, the purine-binding loop (residues 126-134) in the nucleotide-free reconstruction is 3.5Å away from the position found in the Mg^2+^**·**ADP conformation. This shift causes the residue Y134 to point away from the nucleotide-binding site, where it can no longer coordinate the nucleotide purine ring. Residues Y128 and P127, which make hydrophobic contact with the adenine ring, are also shifted away from their ADP-bound positions in the nucleotide-free conformation, prohibiting any significant interaction with the ADP. Furthermore, the loop spanning residues 673-686, is shifted 2.5Å away from its position in the nucleotide-bound form. In total, comparing the active site of the ADP-bound form to the rigor form shows that in the rigor conformation, the active site opens. Here, the residues coordinating the nucleotide are positioned away from the bound nucleotide, which in turn, would promote the release of Mg^2+^**·**ADP. Although we note the structural differences associated with nucleotide release, whether or not ADP release from the active site precedes or succeeds the conformational rearrangement of the active site is not revealed by our methods.

While it appears that the actin-binding regions and ADP-binding site of different myosin isoforms are generally conserved, the intradomain positioning of the converter domain and lever arm differ substantially between myosin species and between the nucleotide-free and ADP-bound states(Pospich et al., 2021; Mentes et al., 2018; Gong et al., 2021, 15). It is these changes that are associated with structural adaptations that allow myosin motors to be kinetically tuned for their respective roles(Nyitrai and Geeves, 2004).

### Small changes in the active site are amplified to large scale structural rearrangements during ADP release

In addition to the differences at the active site, there are comparatively large-scale reorganizations in the lever arm, converter domain, relay helix, and N-terminus when comparing the ADP-bound and rigor structures. Our results provide descriptions of the larger motions accompanying ADP release and point to disease-causing mutations that may disrupt the conformational changes described below.

When comparing the two structures, the most obvious difference is observed at the lever arm. This is characterized by a ∼16° pivot rotation of the helix around residue G768 (see Figure 4B), which results in a ∼16 Å displacement of the most distal residue in the model (essential light chain residue Glu 61). This value is consistent with the displacement associated with the ADP release step of the full-length human cardiac myosin S1, which has been shown to be 15 +/-3 Å as measured by single molecule experiments (Woody et al., 2018). The extent of the lever arm rotation during the ADP release step is larger than that observed for myosin V (9°) but smaller than the shift seen in myosin 1b or smooth muscle myosin (∼25°) (Pospich et al., 2021; Mentes et al., 2018; Volkmann et al., 2000). Importantly, the magnitude of the lever arm swing and the associated changes during ADP release are correlated with the myosin isoform’s degree of force-sensitivity(Greenberg et al., 2016). Under load, when ADP release may be stalled, larger conformational changes between the ADP-bound and rigor states may interfere with such isomerization, thus contributing to a larger force-sensitivity. Accordingly, myosin 1b and smooth muscle myosin, characterized as significant force-sensitive motors, are associated with the largest conformational change, while β-cardiac myosin II is modestly force-sensitive and myosin V as even less sensitive.

Associated with the conformational change of the lever arm, is a change in the converter domain (residues 709-768), here comprised of a β-turn and two helices that are directly adjacent to the lever arm helix. The converter domain sits between the lever arm and the active site, amplifying the small conformational changes in the nucleotide pocket to the large-scale changes of the lever arm and essential light chain. As shown in Figure 4B, comparisons of our ADP-bound and rigor complexes at their actin-binding regions reveals a ∼6.5 Å rigid body shift of the domain (when measured from the Cα atoms of residue 744). In the ADP-bound form, the converter domain is also positioned directly adjacent to the essential light chain, forming an interacting face between light chain residues 123-135 and converter domain residues 720-728. In the rigor form, however, the 6.5 Å shift appears to prevent significant interaction between the two domains. In recent studies, the converter domain has been identified as a hot spot for mutations that are associated with cardiomyopathies, as shown in Figure 5 (Gunther et al., 2019). In fact, recent analysis of the converter domain mutation, F764L, linked to dilated cardiomyopathy, has shown that that the mutation slows myosin ADP release and alters the force-dependent properties of the motor(Tang et al., 2019, 2021). These observations indicate that this region is integral to the mechanochemical coupling of myosin, particularly for the ADP release step.

**Figure 5:**
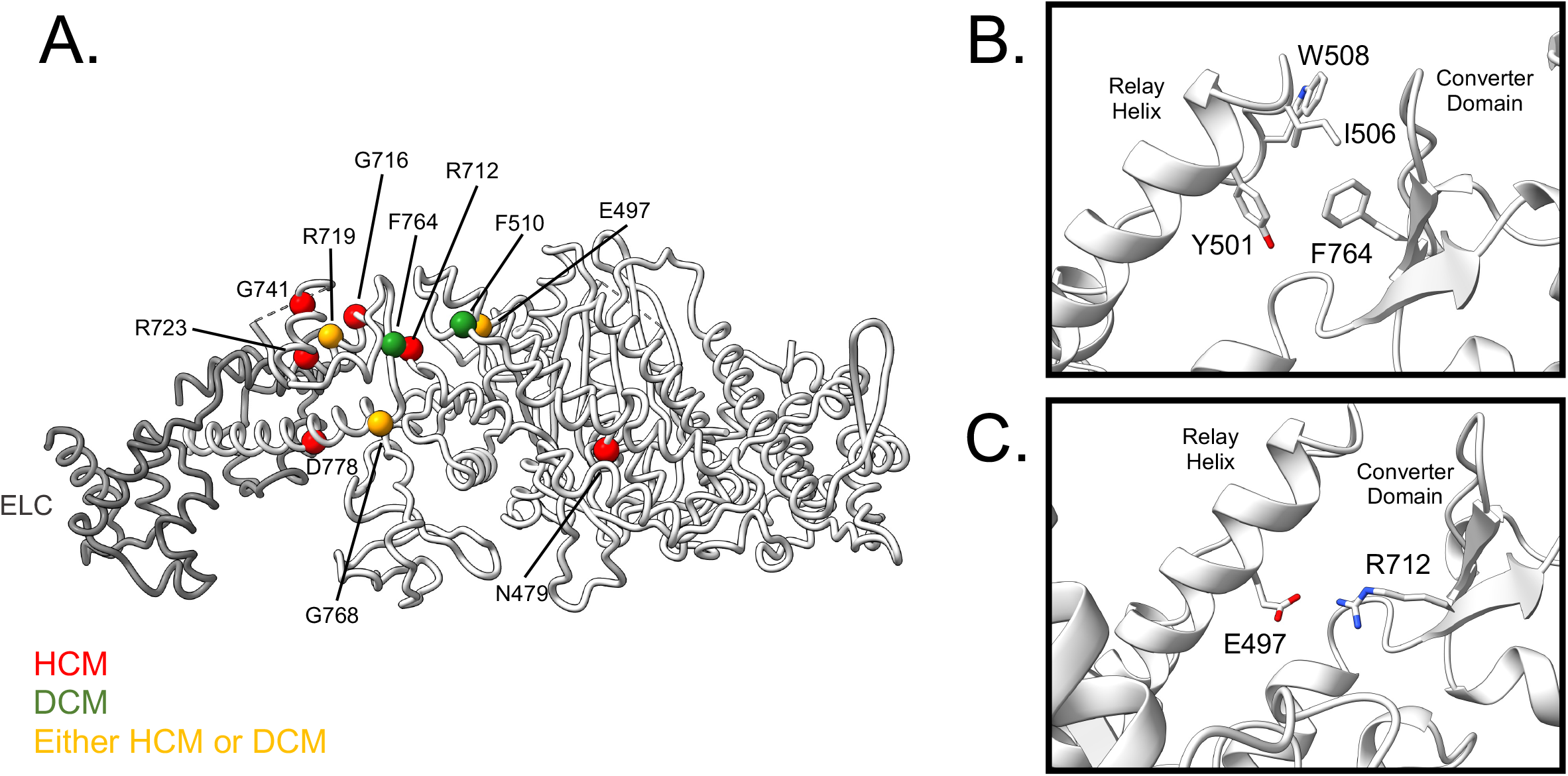
Cardiomyopathy mutations occur in regions involved in ADP release. A). Cartoon diagram of the nucleotide-free β-cardiac myosin II S1 domain highlighting the α-carbon of residues that are known cardiomyopathy-causing mutations. Spheres indicate loci for mutations leading to hypertrophic cardiomyopathy (red), dilated cardiomyopathy (green) or both HCM and DCM (yellow). Each residue was found using the ClinVar archive filtering residues that are known to be pathogenic. B). Shown is the location of myosin residue F764, which sits at the interface between the converter domain and the relay helix. The mutation F764L is known to lead to DCM. C). Shown is the location of the residue R712, which also sits at the interface between the converter domain and the relay helix. Mutations at this site are known to cause HCM. A cluster of mutations are fund in the converter domain, making it a hotspot of mutations.

In addition to these large-scale domain movements, there are several other changes within the motor domain that couple active site rearrangement to force-producing movements. One of these changes occurs in the relay helix. This helix, which is comprised of residues 473-504, traverses the upper 50 kDa domain connecting the active site to the converter domain. Comparing the ADP-bound complex to the rigor form at this region reveals that the edge of the relay helix is shifted nearly 3Å towards the body of the myosin motor, in the same direction of the converter domain associated changes (as measured by the last residues on the relay helix, E504). It appears that the two regions maintain their attachment to each other in both the ADP-bound and nucleotide-free states. The connection of these two regions is facilitated by electrostatic and hydrophobic contacts, including the salt-bridge extending from R712 of the converter to E497 of the relay helix and the hydrophobic pocket created from F764 of the converter and Y501, I506, and W508 of the relay helix (see Figure 5). These conformational changes suggest that the two regions move in concert with each other during nucleotide release, supporting the idea that the relay helix acts as an important link between the active site and converter.

We also observed changes in the N-terminal regions of myosin (residues 5-36). This region connects the active site to the lever arm and thus may play a role in ADP release. In both the ADP-bound and rigor states, residues 8-14 of the N-terminus makes associations with the active site at the purine-binding loop, particularly from the N-terminal residues F9 and the purine-binding loop residue, Y142. While residues 26-36 of the structure extend towards the lever arm in both structures, in the rigor form the lever arm moves ∼8Å towards the N-terminal region, creating an interacting surface that stabilizes the nucleotide-free conformation of the lever arm.

Overall, our structures describe the differences of the human β-cardiac myosin II motor on actin in the presence and absence of ADP. Since the ADP-bound intermediate precedes the rigor conformation in the myosin kinetic cycle, the results presented provide fundamental insights into the conformational changes of the motor during ADP release. The structures we present in this paper show key residue interactions that link changes associated with ADP release in the active site, such as the displacement of the HF helix, purine-binding loop, and the transducer that can be traced to larger scale changes in the lever arm though the transducer, relay helix, converter, and N-terminus of myosin.

## Conclusions

We used cryo-EM and image reconstruction to solve the structures of human cardiac myosin S1 strongly bound to actin-tropomyosin filaments in nucleotide-free and ADP-saturated conformations. We reconstructed human β-cardiac myosin II heavy chain in our work to replicate as faithfully as currently possible the conformations occurring within the human ventricle. Comparison of our structures provides a framework to interpret the functional importance of the ADP release step during the crossbridge cycle and in particular a 16° swing of the lever arm as well as a 6.5 Å movement of the converter domain. Neither of these changes have been previously identified in reconstructions of striated muscle proteins. The lever arm movement between the two myosin states corresponds to a displacement of ∼16 Å and appears to be smaller than that shown for smooth muscle but occurs in the same direction, which is consistent with the corresponding secondary powerstroke steps also exhibited by myosin V, and 1b following ADP release(Mentes et al., 2018; Pospich et al., 2021; Volkmann et al., 2000).

While all myosin motors studied to date (Mentes et al., 2018; Gong et al., 2021; Pospich et al., 2021) display similar conformational interactions in their nucleotide binding core, interactions at the more peripheral converter and lever arm are less conserved, suggesting unique ADP release mechanisms present in different myosins. These distinctions may be associated with differences in force-sensing mechanisms postulated for different myosin isoforms, where, as mentioned, large changes correlate with a higher degree of force-sensitivity (Greenberg et al., 2016), although it remains uncertain how protein sequence and corresponding domain differences contribute. For example, in both myosin 1b and the human β-cardiac myosin II, interactions between the light chain and the motor head, and in particular the converter domain, may stabilize the ADP-bound “up” lever arm position, while the N-terminus of the motor stabilizes the “down” position in the rigor form. In contrast, myosin V lacks these interactions, which may lead to a smaller difference in lever arm angle between the two states. Apart from these structural features, other factors, such as the relative flexibility and orientation of these regions may contribute to differences in ADP release rearrangements.

### Perspective

Cardiac performance is highly regulated and finely tuned to meet physiological demands. Cardiac contractility is not simply controlled by Ca^2+^ levels influencing troponin and tropomyosin. In fact, cardiac sarcomeric proteins are particularly responsive to post-translational chemical modification such as phosphorylation, acetylation and citrullination as well as to mechanical cues such as length change and strain(Hegyi et al., 2021), which in turn modulate cardiac muscle force and power production(Greenberg et al., 2014). For example, increased load applied to cardiac muscle preparations slows ADP release from myosin, and consequently ATP-associated release of myosin from actin to initiate a new crossbridge cycle. The extended lifetime of force-producing myosin on actin would be expected to diminish contraction velocity but increase power output, an adaption thought to affect the motor’s ability to generate power against the afterload. Indeed, in vitro work indicates that counter-balancing forces opposing β-cardiac myosin movement reduces the rate of ADP release from myosin(Greenberg et al., 2014). It follows that load restraining motor head movement on actin in turn will restrict the converter domain and lever arm rearrangements, trapping ADP in the active site, thus providing a likely mechanism for force maintenance(Mentes et al., 2018). In the current study, we have provided structural evidence that this hypothesis can be extended to cardiac muscle.

Our studies suggest that in the presence of an opposing load, the 16° rotation of the lever arm and the 6.5 Å movement of the converter would be restricted in cardiac muscle, mechanically trapping the Mg^2+^·ADP in the active site. By doing so, myosin would tend to become stalled in a strong actin-binding, force-producing state, thereby giving the muscle a mechanism for tuning its total power output. Biochemical studies have shown that disease-associated mutations in myosin that interfere with ADP release can result in severe cardiac dysfunction leading to either hypertrophic or dilated cardiomyopathy(Tang et al., 2021; Snoberger et al., 2021). In fact, many of these mutations are found within the regions of the myosin associated with ADP release and intra-domain movement. For example, in Figure 5, we display the locations of multiple mutations found in the relay helix, converter domain, and lever arm. Indeed, several of these mutations, including F764L and R712L have been shown to disrupt the ADP release step from myosin(Tang et al., 2019, 2021; Snoberger et al., 2021). It is likely that some of these mutations disrupt the proper communication networks that link active site differences to these outer myosin regions. Future structural studies should reveal these connections more explicitly.

## Author Contributions

M.H.D., W.L., and M.J.R. developed the rationale for the project. D.S. and S.M.L.B. generated plasmids, expressed and purified the myosin S1 protein under the direction of C.M.Y. M.E.B. and J.R.M. expressed and purified the actin and tropomyosin protein. M.H.D. carried out all the experimental work described. He assembled the complex, screened and optimized vitrification procedures, and processed all cryo-EM data with help from E.B. M.H.D. built the atomic models with help from M.J.R. M.H.D. interpreted the data, prepared figures, and wrote the manuscript with W.L. and M.J.R.

## Acknowledgements

This project is supported by the NIH grants R01HL036153 to W.L. and R01HL127699-03 to C.M.Y. M.H.D. was supported by NIH Training Program Grant T32HL007969 (to Katya Ravid) and by the Boston University Division of Graduate Medical Sciences institutional funds. Preliminary screening of electron microscope samples was carried out in house and supported by NIH Grant S10RR25434 to E.B.. Final cryo-EM data collections were carried out by the Purdue Cryoelectron Microscopy Facility supported by the NIH Common Fund Transformative High-Resolution Cryoelectron Microscopy Program (U24 GM129541). We would like to thank Thomas Klose and Frank Vago for the collection of the data. Computational work was carried out in house and using resources provided by the Massachusetts Green High Performance Computing Center. The content of the work presented is solely the responsibility of the authors and does not represent the official views of the NIH.

## Data Availability

The structures generated from this work have been deposited in the Protein Data Bank and Electron Microscopy Data Bank under the accession codes: 8EFI, EMD-28083 (helically averaged complex containing ADP), 8EFH, EMD-28082 (helically averaged complex in the rigor conformation), 8EFE, EMD-28081 (focused refinement of the complex containing ADP), 8EFD, EMD-28080 (focused refinement of the complex in the rigor form).

---

**Supporting Figure 1:**
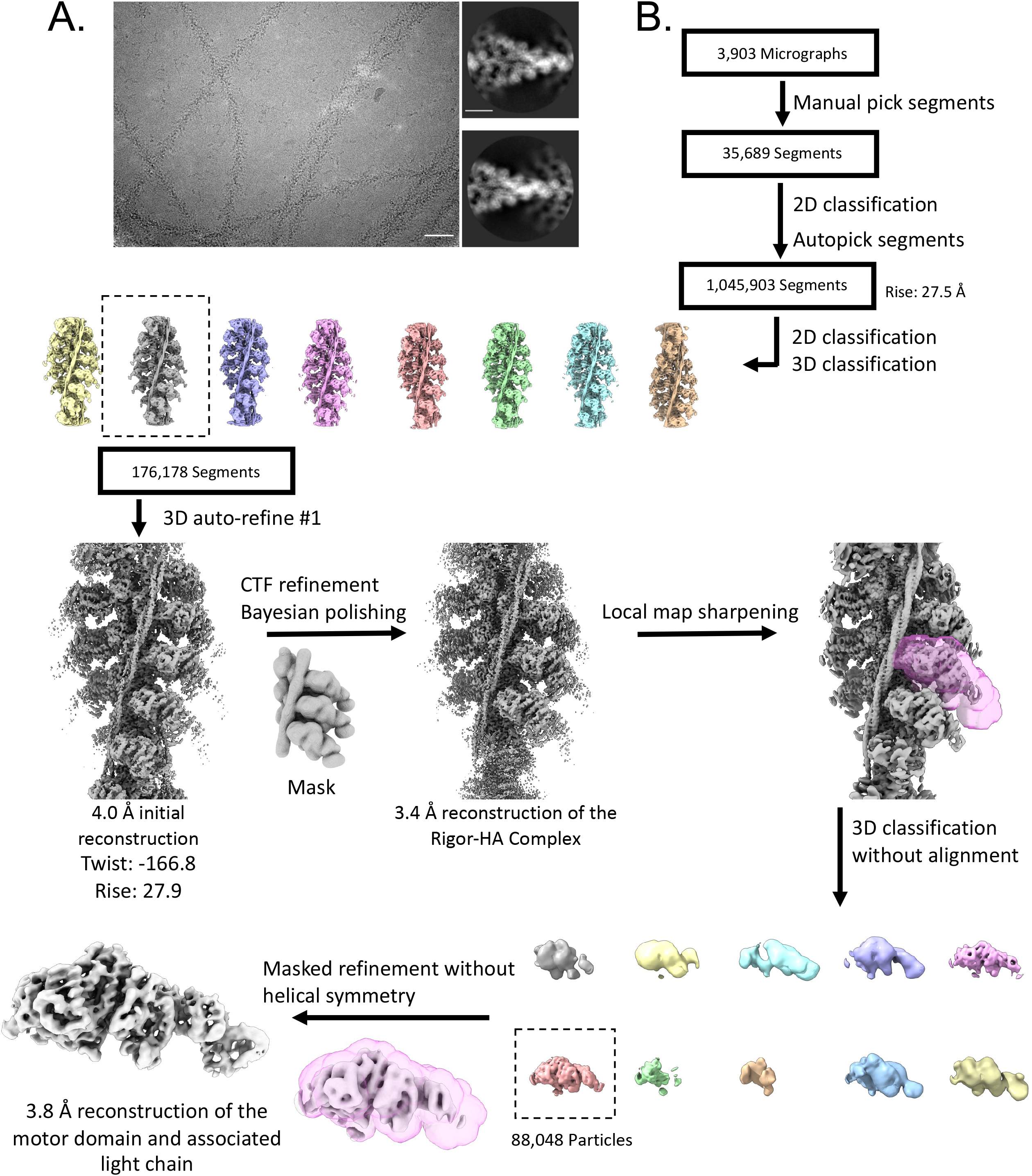
Cryo-EM Image Processing for the Rigor Complex. A). Representative cryo-EM micrograph of the human cardiac actin-tropomyosin-myosin rigor complex, taken from a dataset of 3903. Scale bar: 50 nm. Adjacent to the micrograph are representative 2D class averages of the complex, where the actin, tropomyosin and myosin can be seen in more detail. Scale bar: 5 nm. B). Flowchart describing the cryo-EM data processing scheme to determine the helically averaged human cardiac actin-tropomyosin-myosin rigor complex. This flowchart also includes the focused classification scheme that improved the reconstruction of the myosin-essential light chain complex. Briefly, after generating 2D class averages from manually picked particles, autopicking produced 1,045,903 segments. Subsequent 2D and 3D classifications identified the best segments for use in 3D auto-refine. Further refinements produced a final helically averaged map to 3.4 Å resolution. Focused classification produced the final 3.8 Å reconstruction of the myosin-essential light chain complex.

**Supporting Figure 2:**
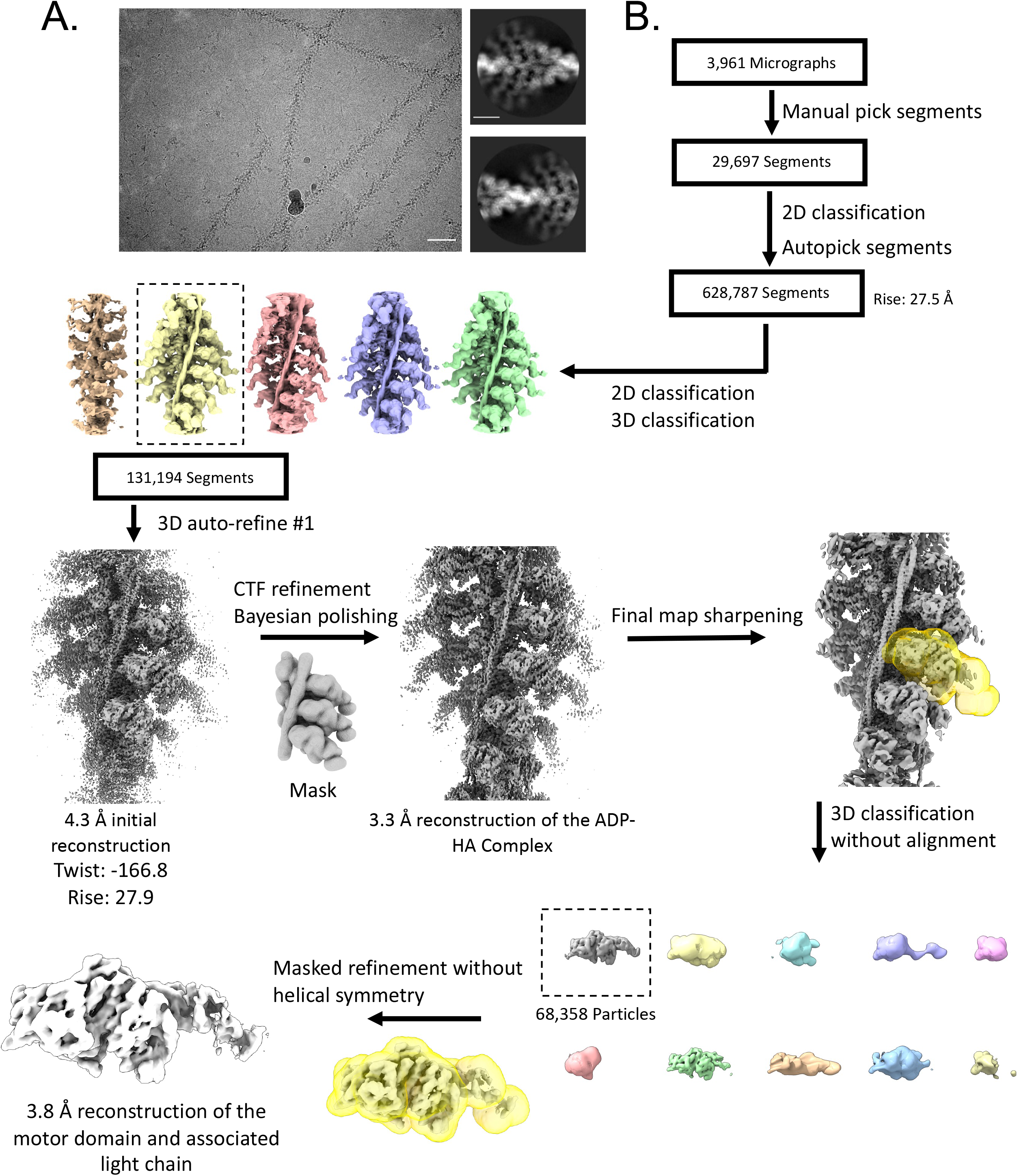
Cryo-EM Image Processing for ADP-Mg^2+^ Complex. A). Representative cryo-EM micrograph of the human cardiac actin-tropomyosin-myosin complex bound to ADP-Mg^2+^, taken from a dataset of 3961. Scale bar: 50 nm. Adjacent to the micrograph are representative 2D class averages of the complex, where the actin, tropomyosin and myosin can be seen in more detail. Scale bar: 5 nm. B). Flowchart describing the cryo-EM data processing scheme to determine the helically averaged human cardiac actin-tropomyosin-myosin complex bound to ADP. This flowchart also includes the focused classification scheme that improved the reconstruction of the myosin-essential light chain complex. Briefly, after generating 2D class averages from manually picked particles, autopicking produced 628,787 segments. Subsequent 2D and 3D classifications identified the best segments for use in 3D auto-refine. Further refinements produced a final helically averaged map to 3.3 Å resolution. Focused classification produced the final 3.8 Å reconstruction of the myosin-essential light chain complex bound to ADP.

**Supporting Figure 3:**
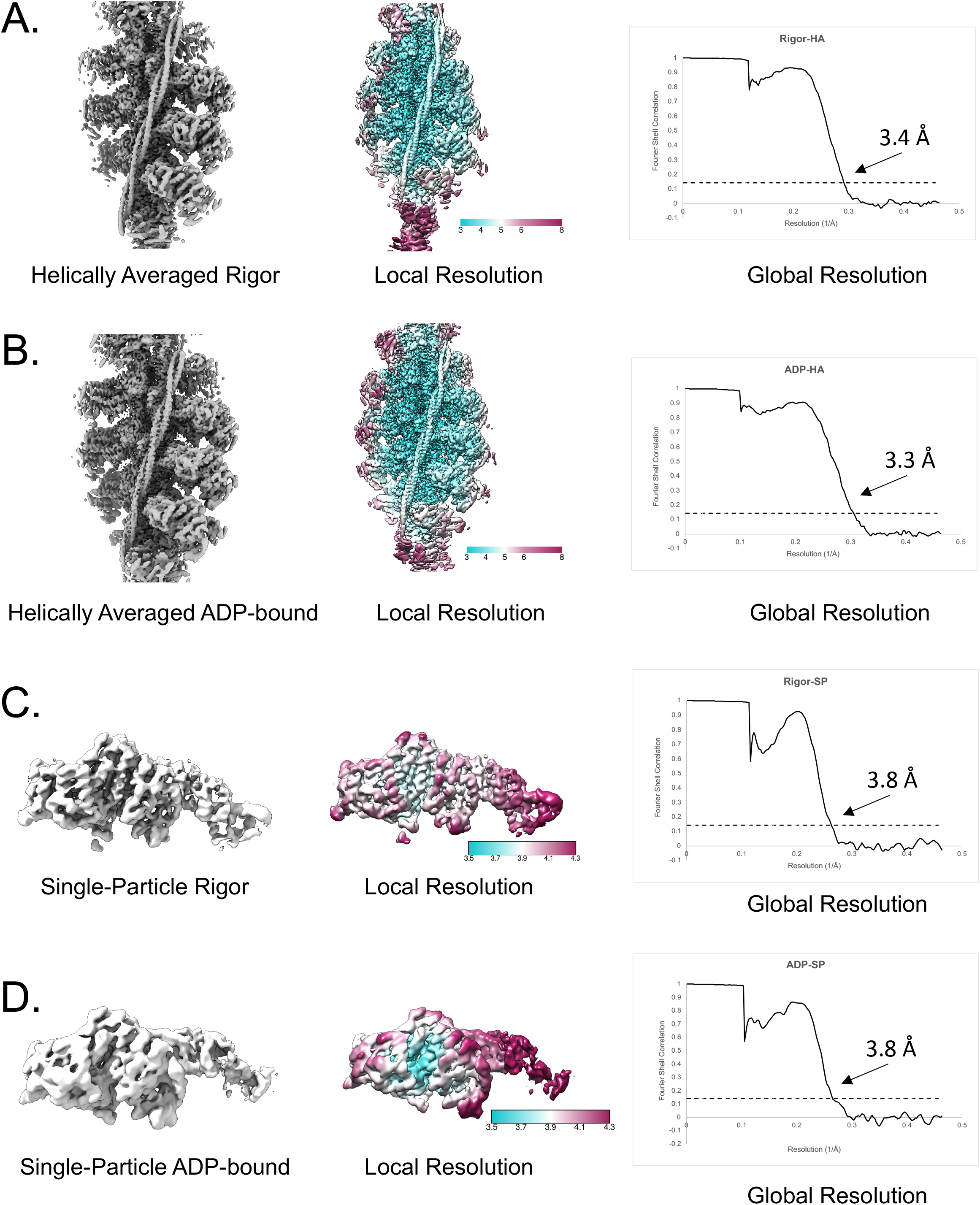
Maps and FSC Curves. A). Final map, local resolution distribution, and final FSC curve for the helically averaged rigor complex. The local resolution was calculated in RELION. The same is shown for the helically averaged ADP-bound complex (B), the focused refinement for the rigor form (C), and the focused refinement for the ADP-bound complex (D).

**Supporting Figure 4:**
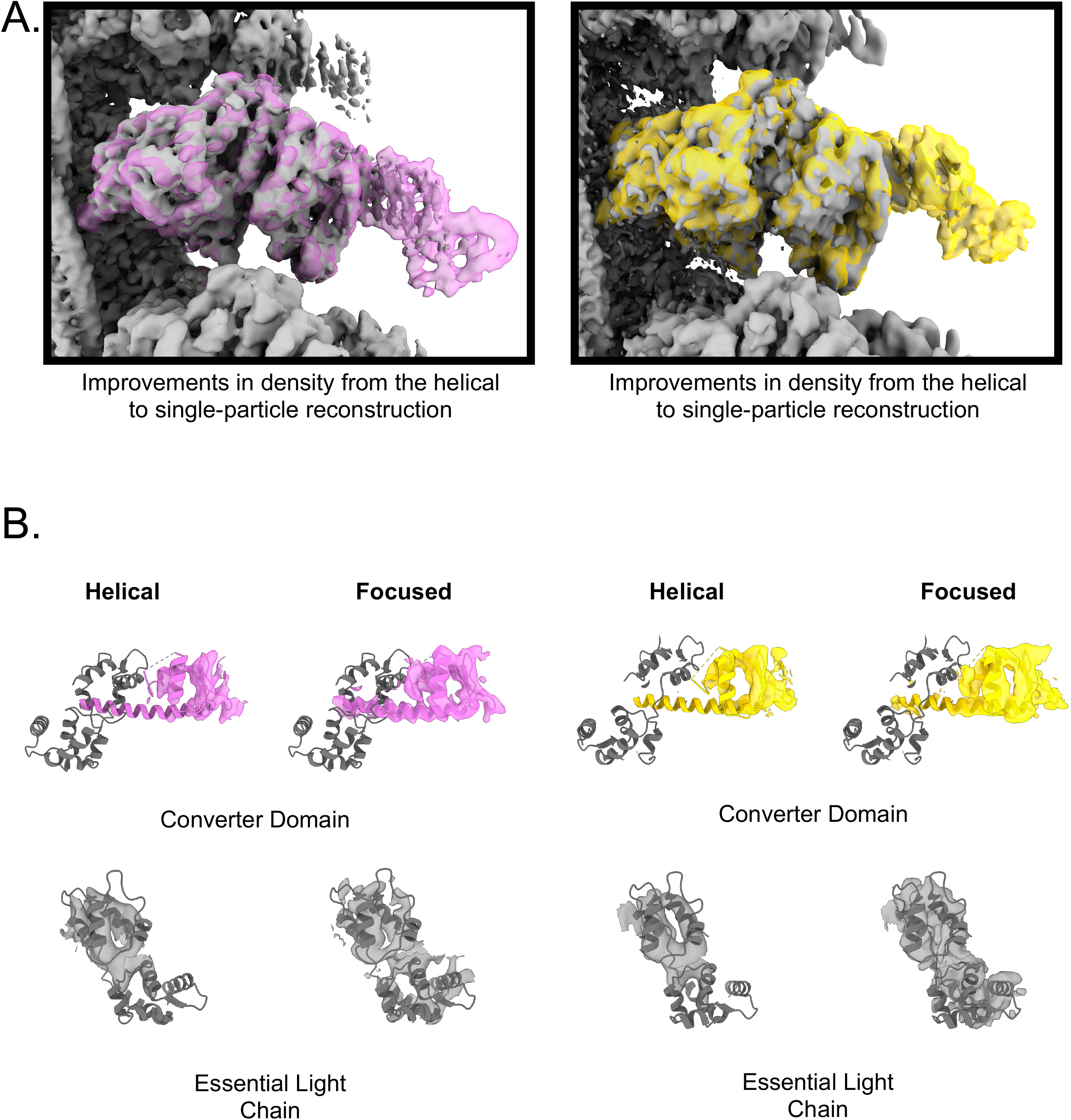
Improvements from Focused Refinements. A). Improvements in density from the helical to single-particle reconstruction for the rigor and ADP-bound complexes. The helically averaged reconstruction for both complexes is shown in grey, while the focused refinements are overlayed colored orchid (rigor) and yellow (ADP-bound). B). Improvements to specific regions of the motor domain, including the lever arm and converter as well as the essential light chain.

